# TRPC6 is a mechanosensitive channel essential for ultrasound neuromodulation in mammalian brain

**DOI:** 10.1101/2024.03.06.583779

**Authors:** Yumi Matsushita, Kaede Yoshida, Miyuki Yoshiya, Takahiro Shimizu, Satoshi Tsukamoto, Yuichi Takeuchi, Makoto Higuchi, Masafumi Shimojo

## Abstract

Ultrasound neuromodulation has become an innovative technology that enables non-invasive intervention in mammalian brain circuits with high spatiotemporal precision. Despite the expanding utility of ultrasound neuromodulation in the neuroscience research field and clinical applications, the molecular and cellular mechanisms by which ultrasound impacts neural activity in the brain are still largely unknown. Here, we report that transient receptor potential canonical 6 (TRPC6), a mechanosensitive non-selective cation channel, is essential for ultrasound neuromodulation of mammalian neurons *in vitro* and *in vivo*. We first demonstrated that ultrasound irradiation elicited rapid and robust Ca^2+^ transients mediated via extracellular Ca^2+^ influx in cultured mouse cortical and hippocampal neurons. Ultrasound-induced neuronal responses were massively diminished by blocking either the generation of action potential or synaptic transmission. Importantly, both pharmacological inhibition and genetic deficiency of TRPC6 almost completely abolished neuronal responses to ultrasound. Furthermore, we found that intracerebroventricular administration of a TRPC6 blocker significantly attenuated the population of neuronal firings in the cerebral cortex evoked by transcranial ultrasound irradiation in mice. Our findings indicate that TRPC6 is an indispensable molecule of ultrasound neuromodulation in the intact mammalian brains, providing fundamental understanding of biophysical molecular mechanisms of ultrasound neuromodulation as well as insight into its future feasibility in neuroscience and translational researches in humans.

## Introduction

Neuromodulation is a cardinal technique for altering neuronal activity in the brain circuit by delivering various external stimuli such as electrical currents, magnetic field, and acoustic irradiation. These approaches enable non-invasive, non-surgical intervention in the neuronal activity of targeted tissue to understand brain function in basic neuroscience researches, and to therapeutically ameliorate brain dysfunctions in neurological disorders (1, 2). For instance, transcranial direct current stimulation (tDCS) and transcranial magnetic stimulation (TMS) have recently been established as neuromodulation devices that can alter the neocortical activity of the human brain, and have accomplished considerable benefit in therapeutical applications for epilepsy, depression, and disorders of consciousness (3). However, these conventional modalities affect a relatively broad range of the brain surface and are not suitable for regulating neuronal activity in a limited area within a millimeter scale or a deep brain region (4, 5). In contrast, ultrasound is currently attracting great attention as an innovative technology that may change the basic principle of neuromodulation. Although the potential applicability of ultrasound for neuromodulation had already been reported in the 1920s (6), equipment performance was insufficient for generating adequate irradiation power for transmission of the skull, thereby restricting its brain application during the last decades (7). Nonetheless, recent technical advances in focused acoustic waves have overcome most of the limitations; ultrasound has received renewed interest as a versatile tool for noninvasively controlling brain circuits with excellent tissue penetration and high spatial precision (8, 9).

Despite the expanding utility of ultrasound in the mammalian brain, the mechanisms by which ultrasound alters brain function remain a fundamentally remain an open question. In rodent brains, transcranial ultrasound may directly impact neuronal activity *in vivo* either by stimulation of targeted tissue (10, 11) or by indirect effect via the mechanism related to the auditory system (12, 13). Therefore, identification of the key components involved in the mechanism of ultrasound neuromodulation is difficult by relying only on *in vivo* experimental systems. On the other hand, mouse cortical neurons in dissociated cultures and brain slices lacking any auditory influences also demonstrate robust responses to ultrasound stimulation (14–16). These facts indicate that neurons have an intrinsic ability to directly respond to ultrasound. Given the major bioeffects of ultrasound in living organisms such as mechanical pressure and cavitation (9), these findings indicate that innate mechanosensitive molecules in neurons are critically involved in ultrasound neuromodulation. In fact, mammalian neurons express various types of mechanosensitive ion channels that sense biophysical changes both inside and outside the body (17), and some of them have already been pointed out as essential players in ultrasound neuromodulation (5). Nevertheless, experimental evidence provided by previous studies is still controversial and the crucial mechanosensitive channel that contributes to ultrasound neuromodulation has not yet been identified. These issues prompted us to further explore the molecular and cellular bases of the ultrasound-mediated regulation of neuronal functions.

In this research, we established a live-cell imaging assay system based on the usage of a fluorescence microscope combined with an ultrasound stimulator and systematically characterized ultrasound-elicited cellular responses in cultured cortical and hippocampal neurons. In parallel, using the same technical paradigm of ultrasound stimulation, we also efficiently validated whether the findings obtained from cell culture-based analysis were consistent with the observation from living mouse brains by *in vivo* electrophysiological recordings. Notably, our strategy identified transient receptor potential canonical 6 (TRPC6), a mechanosensitive and non-selective cation channel, as a key sensor molecule responsible for the initial biological event of intrinsic neuronal response to ultrasound neuromodulation in mammalian neurons both *in vitro* and *in vivo*.

## Materials and methods

### Animals

Animal experiments were performed and reported according to the National Research Council’s Guide for the Care and Use of Laboratory Animals, institutional guidelines of the National Institutes for Quantum Science and Technology and Hokkaido University, and the Animal Research: Reporting in Vivo Experiments (ARRIVE) guidelines. C57BL/6J mice were purchased from Japan SLC (Hamamatsu, Japan) or Oriental Yeast Co., Ltd. (Itabashi, Tokyo). TRPC6 deficient mouse strain (TRPC6-KO; B6;129S-*Trpc6^tm1Lbi^/*Mmjax, Stock # 37345-JAX) was purchased from the Jackson Laboratory (Bar Harbor, ME, USA) (18). All mice were housed under specific-pathogen free condition and maintained in a 12 h light/dark cycle with ad libitum access to standard diet and water.

### Cultured neuron preparation *in vitro*

Primary neuronal culture was prepared as described previously (19). Briefly, neocortex and hippocampus of brain tissues were dissected from either C57BL/6J or TRPC6-KO mice at embryonic days 16 to 18. Neurons were isolated by digesting the collected tissues using papain (Worthington, Lakewood, NJ, USA) in Hank’s Balanced Salt Solution (HBSS, Invitrogen, Waltham, MA, USA). The isolated cells suspended in Neurobasal-A medium (Invitrogen) supplemented with 10% fetal bovine serum (HyClone, Cytiva, Tokyo, Japan), 1% GlutaMAX (Invitrogen), and 1% N2 supplement (Invitrogen), and plated onto Poly-D-Lysine coated 12 mm glass coverslips (Corning, NY, USA) at a density of 250,000 - 300,000 cells/cm^2^. The next day (DIV1), the medium was replaced by Neurobasal-A medium supplemented with 2% fetal bovine serum, 1% GlutaMAX, and 2% B27 supplement (Invitrogen). For calcium imaging, neuronal cells at DIV1 were infected with a recombinant adeno-associated virus (AAV) encoding GCaMP6s, a genetically encoded fluorescent calcium indicator, driven by synapsin promoter (Addgene, 100843-AAV9, 2×10^9^ vg/coverslip) (20). The neurons were maintained by replacing half of the medium with fresh Neurobasal-A medium supplemented with 1% GlutaMAX and 2% B27 supplement every 3 days and used for experiments at DIV12 to DIV16.

### Ultrasound and electrical stimulation *in vitro*

Ultrasound stimulation was conducted using a 6-mm diameter flat transducer (ST-TM1-6, NEPA GENE, Chiba, Japan) driven by SonoPore KTAC-4000 (NEPA GENE), an ultrasound amplifier to provide programmed burst pulses. For acoustic stimulation of the neuronal culture, a tip of the transducer was immersed in a bath solution and placed 10 mm above neurons on the coverslip in the glass-bottom imaging chamber (CK-1, Narishige, Tokyo, Japan), and ultrasound irradiation was directly applied to the neurons via the bath solution. Correlation between the distance from the tip of the transducer and acoustic intensity was measured with an ultrasound power meter (UPM-DT-10PA, OHMIC Instruments, St. Charles, MO, USA), and we confirmed no significant difference in intensity within a range of 5 mm to 28 mm (less than 15%). We used 1-MHz ultrasound burst pulses of 100 Hz in pulse repetition frequency and 50% in duty cycle (Fig. 1B) as otherwise described. For electrically-evoked field stimulation of neuronal culture, a tip of tungsten concentric bipolar microelectrode (World Precision Instruments, Sarasota, FL, USA) was positioned 1.5 µm above the neurons. To induce neuronal excitation, the extracellular electric train stimulus composed of 10 pulses (each pulse given at 10 V at a duration of 1 msec) was generated by Master-8 pulse stimulator with ISO-FLEX flexible stimulus isolator (A.M.P.I., Jerusalem, Israel).

**Figure 1.**
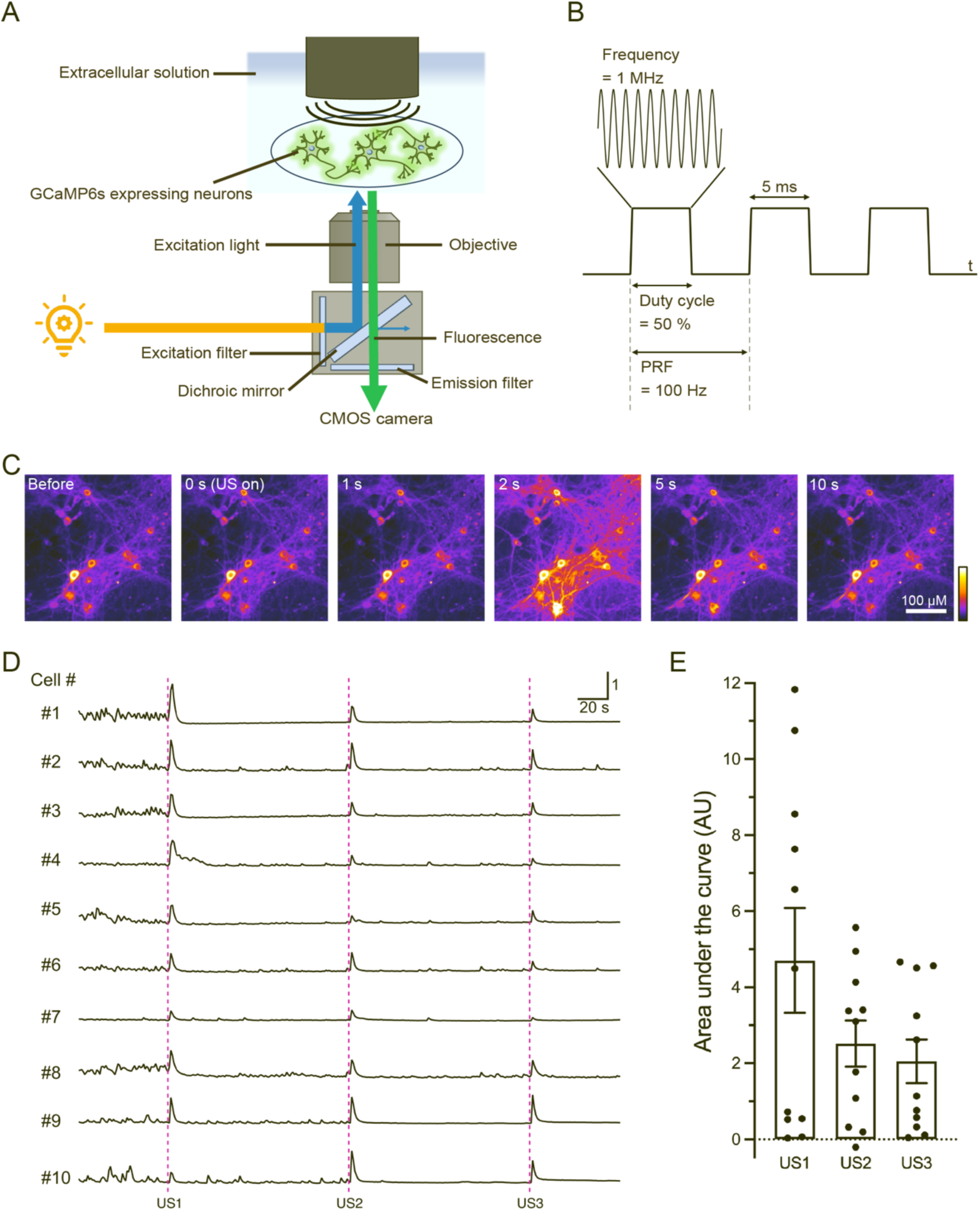
Ultrasound irradiation induces Ca^2+^ transients in cultured cortical neurons. A) A scheme of *in vitro* experimental setup to monitor ultrasound-induced neuronal response. B) A schematic illustration of the acoustic waveform and ultrasound stimulation parameters used in this study. PRF: Pulse repetition frequency. C) Representative sequential images depicting GCaMP6s fluorescence in neuronal cells during pre-, post-, and peri-stimulation phases induced by ultrasound irradiation. D) Representative traces of fluorescence calcium imaging demonstrating pulsed ultrasound irradiation induced rapid Ca^2+^ transients in individual neurons. Ultrasound stimulation at the intensity of 190 mW/cm^2^ and duration of 500 msec is applied to neurons at the time of US1, US2, and US3. E) The amplitudes of Ca^2+^ transients in responses to sequential ultrasound stimulation at US1, US2, and US3 are quantified by area under the curve (AU). Bar graph values show mean ± SEM (n = 11, *P* = 0.1349, one-way ANOVA).

### Fluorescence calcium imaging

Fluorescence calcium imaging for optical recording of neuronal activity was performed with an inverted widefield fluorescence microscope (IX83, Olympus, Tokyo, Japan) controlled by Olympus Cell Sense Dimension software. Cultured neurons expressing GCaMP6s grown onto a coverslip were transferred to the imaging chamber attached to the stage of the microscope. During each imaging session, neurons were maintained by continuous perfusion with HEPES-buffered bath solution (130 mM NaCl, 5 mM KCl, 10 mM HEPES, 10 mM D-glucose, 2 mM CaCl_2_, 1 mM MgCl_2_) at a rate of 1.5–2 mL/min. For time-series analyses of changes in fluorescence intensity, high-resolution images were captured at 1 Hz by Orca-Flash4.0 CMOS camera (Hamamatsu Photonics, Hamamatsu, Japan) with x20 objective lens (PlanApo, NA 0.45) at room temperature (22–25°C). Acquired image stacks were next analyzed by NIH ImageJ / Fiji software. To quantify the fluorescence signal intensity of GCaMP6s as an intracellular Ca^2+^ concentration in individual neurons, regions of interest (ROI) including soma of single neurons were manually placed in each image. Approximately 30–60 ROIs per coverslip were randomly selected to measure the Ca^2+^ transients of each neuron, and the averaged grayscale values of each ROI were extracted for data plotting and quantitative analysis. Baseline fluorescence (F_0_) was calculated as the averaged fluorescence intensities for 30 sec during the pre-stimulation period. Normalized fluorescence value (ΔF/F_0_) was then determined as ΔF/F_0_ = (F(t)-F_0_) / F_0_. For measuring the peak magnitude of Ca^2+^ transient elicited by stimuli, the ΔF/F_0_ values for 10 sec during the post-stimulation period were summed to calculate the area under the curve. All data analysis and plots were prepared using Igor Pro (WaveMetrics, Portland, OR, USA) and GraphPad Prism (GraphPad Software). The heatmaps of normalized fluorescence intensity (ΔF/F_0_) of GCaMP6s in neurons were generated using MATLAB (MathWorks, Natick, MA, USA).

### Pharmacological *in vitro* experiments

A Ca^2+^-free solution was prepared by simply omitting CaCl_2_ from the bath solution. Cadmium at 100 µM was used as a nonselective Ca^2+^ ion channel blocker. Tetrodotoxin (TTX; Tocris Bioscience, Bristol, UK) at 1 µM was used to prevent the generation of action potentials. To inhibit synaptic transmission, CNQX (Tocris Bioscience) and AP5 (Tocris Bioscience) were used as selective antagonists against AMPA and NMDA receptors at 10 µM and 50 µM, respectively. Ruthenium Red (Cayman Chemical, Ann Arbor, MI, USA), GsMTx4 (PEPTIDE INSTITUTE, Osaka, Japan), BI-749327 (MedChemExpress, Monmouth Junction, NJ, USA), and Dooku1 (MedChemExpress) were used to block TRPVs channels, mechanosensitive ion channels, TRPC6 channel, and Piezo1 at 5 µM, 10 µM, 1 µM, and 10 µM, respectively. Hyp9 (MedChemExpress) was used to activate TRPC6 at 10 µM. All chemicals described above were added to the bath solution in each experimental session.

### Virus preparation for overexpression

For the overexpression of TRPC6 protein, a cDNA fragment encoding mouse TRPC6 (Accession# NM_013838) fused to HA epitope tag was synthesized and subcloned into the multi-cloning site of a lentivirus shuttle plasmid containing human Ubiquitin C promoter. HEK293T cells were maintained in high-glucose DMEM (Invitrogen) supplemented with 10% fetal bovine serum (Sigma) and Penicillin/Streptomycin (Invitrogen) at 37°C in a 5% CO_2_ incubator. Recombinant lentivirus packaging was performed in HEK293 cells by co-transfection of lentivirus shuttle plasmid and packaging plasmids pVSVg, pRev, and pGag/Pol using FuGENE reagent (Promega, WI, USA). 48 hours after transfection, lentivirus particles secreted in the culture medium were harvested, purified with a 0.45 µm filter, and stored at −80°C until use. The lentivirus was infected to cultured neurons at DIV5 by direct addition of viral solution into the culture medium (30 µl virus per 1 mL).

### RT-PCR

Total RNA was extracted from cultured neurons using RNeasy Mini Kit (Qiagen, Hilden, German) and purified using RNase-Free DNase Set (Qiagen) according to the manufacturer’s instructions. Reverse transcription was performed with 500 ng of total RNA and PrimeScript™ RT Master Mix (Perfect Real Time) (Takara, Tokyo, Japan). Reverse transcription PCR (RT-PCR) was performed with 2 µL cDNA, primers for TRPC6 (from 5’ to 3’: sense; AAA GAT ATC TTC AAA TTC ATG GTC, antisense; CAC GTC CGC ATC ATC CTC AAT TTC), and Q5 High-Fidelity 2X Master Mix (New England Biolabs, Ipswich, MA, USA). RT-PCR protocol was involved in 35 cycles of PCR at 98°C for 10 sec, at 59°C for 30 sec, and at 72°C for 30 sec in TaKaRa Thermal Cyclar Dice Touch (Takara).

### Electrophysiological *in vivo* experiments

Mice were anesthetized with 0.5–1.5% isoflurane followed by subcutaneous administration of 0.1–0.3 mg/kg atropine and mounting on a stereotaxic frame. Ground and reference screw electrodes were fixed on the skull above the cerebellum. A small cranial window (∼1 mm in diameter) was made at 2.06 mm posterior from bregma and 0.36 mm rightward from midline. A tungsten microelectrode (TE0.2-70-B-2; UNIQUE MEDICAL Co., Ltd., Tokyo, Japan) was inserted through the cranial window and advanced at a 45° angle toward the right side of the coronal plane. Signals via the microelectrode were fed to a preamplifier (C3324; Intan Technologies, CA, U.S.A.), and the amplified signals were recorded by recording controller (C3004, Intan Technologies) at a sampling rate of 20 kHz. A 3 mm-diameter flat transducer (ST-T1-3, NEPA GENE) was placed above the right side of the skull and its tip was acoustically coupled to the skull with an ultrasound gel (LOGIQLEAN, GE HealthCare). At each recording site, several recording sessions at different US intensities (0, 70, 130, 210, 350, 520, 730 mW/cm^2^) were performed. One hundred millisecond-long ultrasound irradiations (1 MHz sinusoidal wave, Duty 50%, PRF 100 Hz) were performed 20 times with a random interstimulus interval ranged from 10 to 20 sec. After recordings at each site, the electrode was advanced by 50–100 µm for the next recording sessions.

For pharmacological intervention of TRPC6, BI-749327 (MedChemExpress) was dissolved with a vehicle composed of 10% DMSO, 40% PEG300, 5% Tween-20 and 45% physiological saline, and stored at 4°C at a concentration of 5 mM. Before the experiments, the stock solution was further diluted to 0.6 mM with saline and 2 µl of the diluted solution was intracerebroventricularly administered after the control recording sessions using a NANOFIL micro-syringe (World Precision Instruments; FL, U.S.A.) and ultra-micro pomp (UMP3T-1, World Precision Instruments) at a flow rate of 10 nl/s. Recordings were resumed 30 min after administration.

After the data acquisition, 100 µA anodal direct currents were passed for 10 sec via the electrode tip for the electrical lesion to locate the recording sites. The mice were then transcardially perfused with physiological saline followed by 4% PFA and 0.2% picric acid in 0.1 M PB. After post fixation and cryo-blockade with sucrose infiltration, 50 µm-thick coronal brain sections were prepared with a cryostat. The sections were stained with 1 µg/ml DAPI in 0.1 M PB for 15 min, coverslipped and observed by confocal microscopy (LSM900, Carl Zeiss, Oberkochen, Germany).

### Detection ultrasound-modulated population neural activity *in vivo*

For the detection of modulated neural activity by ultrasound irradiation, peri-stimulus time histograms (PSTH) were prepared for each recording session, and cross-correlation (CCG) analysis was conducted between timestamps of stimulus and population (multi-unit) neural activities with MATLAB (MathWorks, Natick, MA, U.S.A.). Randomly time-jittered 1000 datasets of timestamps of population neural activities were generated by adding random time jitters ranging from −3 sec to +3 sec to the real timestamps of the population neural activities. Pointwise 95% acceptance bands were calculated from the jittered datasets for each 100 msec bin. Multiple comparison error was corrected by introducing ‘global significance bands’ (21). The ultrasound-induced modulation of population neural activities was considered to be significant if any of its CCG bins went across the global significance bands within 1000 msec after the US irradiation.

### Statistics

Statistical analysis was performed by either GraphPad Prism or Microsoft Excel software. Unpaired Student’s t-test and one-way ANOVA followed by Dunnett’s multiple comparisons test were employed for comparisons between two-group data and three- or more group data of *in vitro* experiments, respectively. Χ^2^ test for independence was employed to test the data of fractions of significantly modulated population neural activities of the *in vivo* experiment.

## Results

### Ultrasound irradiation induces an increase of intracellular Ca^2+^ concentration in cultured neurons

To explore the intrinsic mechanisms of neuronal response against ultrasound-mediated neuromodulation, we first set up an experimental system that enables the monitoring of neuronal activity and controlled stimulation by ultrasound waves simultaneously. We employed fluorescence calcium imaging to assess intracellular Ca^2+^ dynamics as an optical indicator of cellular activity. Mouse neurons expressing GCaMP6s, a genetically encoded fluorescence calcium indicator, were grown on a coverslip and placed in an imaging chamber perfused with extracellular bath solution on the stage of an inverted widefield fluorescence microscope. In this configuration, the tip of the ultrasound transducer was dipped into the bath solution and set above neurons in a field of view, and fluorescence time-lapse imaging was conducted by capturing sequential images with a CMOS camera from the bottom side (Fig. 1A). Given the previous studies summarizing the biological applicability of ultrasound-mediated neuromodulation with approximately 0.25 - 2 MHz wave-frequency (22), neurons were stimulated with ultrasound waves at a frequency of 1 MHz. Each stimulus consisted of 100 Hz PRF with a duty cycle of 50% (Fig. 1B).

To establish the appropriate conditions for ultrasound stimulation in which neurons demonstrate a steady response, neurons were sonicated at different stimulation parameters in terms of irradiation intensities (17, 190, 294 mW/cm^2^) and durations (100, 200, 500 msec). At an intensity of 17 mW/cm^2^, neurons barely respond even to a prolonged train of stimulus with a duration of 500 msec. However, ultrasound waves at 190 mW/cm^2^ constantly induced a significant increase in intracellular Ca^2+^ concentration in neurons, and these cellular responses became more robust in a duration-dependent manner (Supplemental Fig. 1). In contrast, at an intensity of more than 294 mW/cm^2^, ultrasound irradiation frequently causes unstable neuronal responses presumably due to the unpredictable mechanical fluctuation and/or biophysical phenomena. Considering these observations, we chose stimulation parameters at an intensity of 190 mW/cm^2^ and a duration of 500 msec in all subsequent experiments. In this optimized protocol, a rapid increase in the fluorescence intensity of GCaMP6s in neurons was induced within a few seconds immediately after the application of ultrasound stimuli, and these neuronal responses were transient and synchronized among neighboring neurons (Fig. 1C). Onset kinetics and the peak amplitude of neuronal responses to ultrasound were also similar to those of neuronal responses elicited by the train of conventional electrical stimuli (Supplemental Fig. 2A). In addition, representative fluorescence traces in independent neurons demonstrate that neurons were repeatedly activated by multiple ultrasound stimulus and neuronal responses varied in magnitude (Fig.1D). Although the data sets of the respective experimental sessions were relatively variable, the average peak amplitude in response to three sequential ultrasound stimuli were in the same range and did not differ significantly (Fig. 1E).

### Neuronal responses to ultrasound depend on extracellular Ca^2+^ influx and network excitation

Since ultrasound stimuli induced rapid Ca^2+^ transients in neurons within a few seconds, the major source of the increased intracellular Ca^2+^ was assumed to be Ca^2+^ influx from outside of the cell. To address this fundamental question, we first investigated whether Ca^2+^ removal from extracellular bath solution impacts ultrasound mediated intracellular Ca^2+^ dynamics in neurons (Fig. 2A). As expected, we observed that extracellular Ca^2+^ free condition drastically suppressed the Ca^2+^ transients even after the ultrasound stimulus (Fig. 2B-D). In addition, ultrasound-mediated neuronal responses were also strongly attenuated in the presence of Cd^2+^, a potent non-selective calcium channel blocker (Fig. 2B-D), suggesting that ultrasound-induced neuronal responses rely on extracellular Ca^2+^ influx through calcium channels on the plasma membrane.

**Figure 2.**
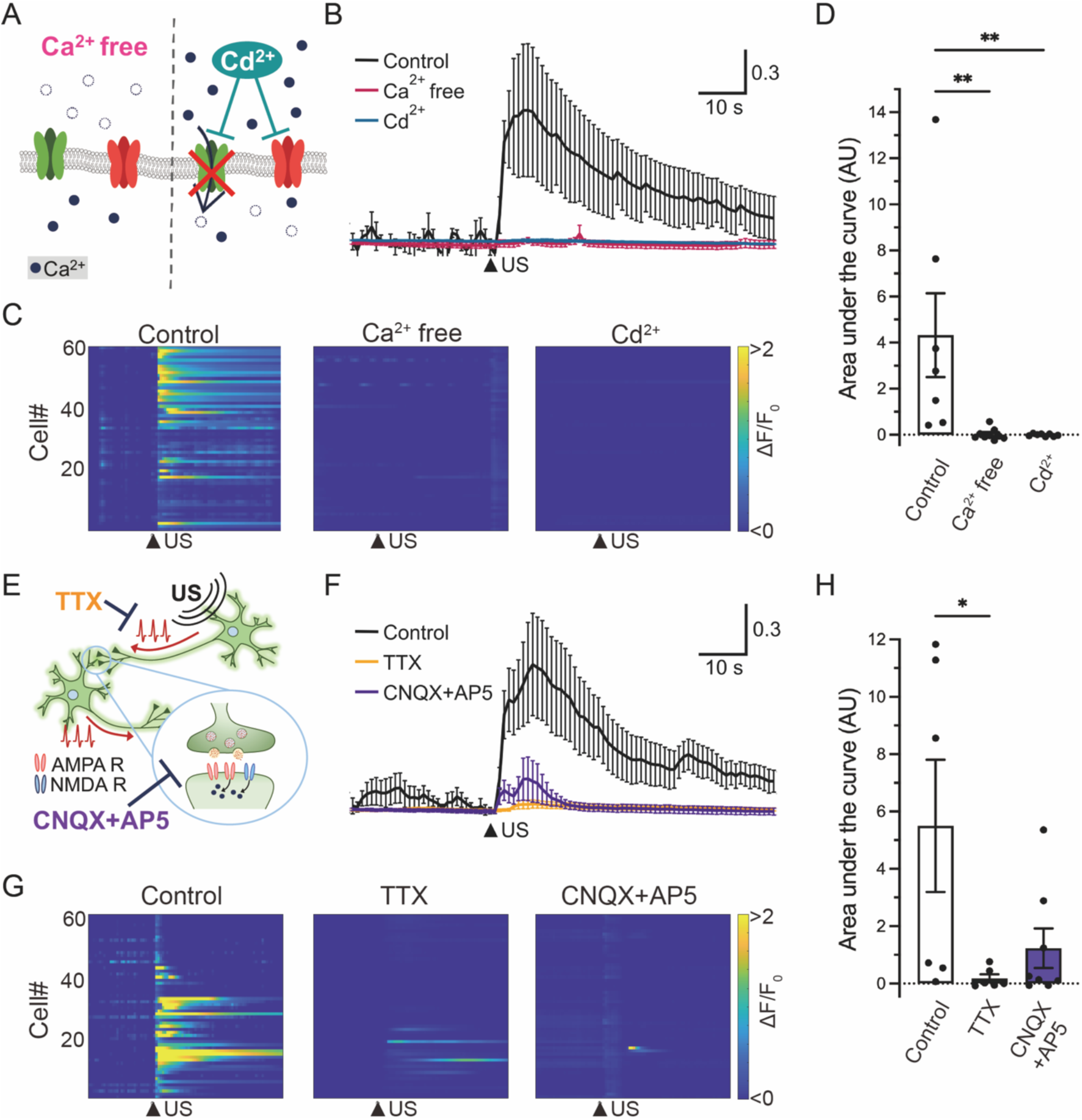
Ultrasound mediated Ca^2+^ transients depend on extracellular Ca^2+^ influx and network activity. A) A scheme of strategies to investigate the impact of Ca^2+^ influx from outside of the cell on ultrasound-induced neuronal response. Cd^2+^ (100 µM) blocks Ca^2+^-permeable ion channels. B) Averaged traces of neuronal Ca^2+^ transients against ultrasound irradiation under the condition of control (black), Ca^2+^ free (red), or the presence of Cd^2+^(blue). Data from seven, nine, and eight independent experiments for control, Ca^2+^ free, and Cd^2+^ are plotted as mean ± SEM, respectively. Arrowhead indicates the time of ultrasound stimulation. C) Heatmap demonstration of normalized fluorescence intensity (ΔF/F_0_) of GCaMP6s in neurons under the condition of control, Ca^2+^ free, or the presence of Cd^2+^. Data from 60 cells in an experiment are plotted. Arrowheads indicate the time of ultrasound stimulation. D) The amplitudes of Ca^2+^ transients in response to ultrasound stimulation are quantified as the area under the curve (AU). Bar graph values show mean ± SEM. \*\**P* < 0.01 (one-way ANOVA followed by Dunnett *post-hoc* test). E) A scheme of strategies to investigate the impact of the generation of action potentials and functional network activity on ultrasound-induced neuronal response. TTX (1 µM) is used to prevent the generation of action potentials and a mixture of CNQX (10 µM) and AP5 (50 µM) is used to block synaptic transmission. F) Averaged traces of neuronal Ca^2+^ transients against ultrasound irradiation under the condition of control (black), TTX-treated (orange), or CNQX+AP5-treated (purple) neurons. Data from six, six, and eight independent experiments for control, TTX, and CNQX+AP5 were plotted as mean ± SEM, respectively. G) Heatmap analysis of neuronal responses under the condition of control, TTX-treated or CNQX+AP5-treated neurons. Data from 60 cells in an experiment were plotted. H) Bar graph values show mean ± SEM. \**P* < 0.05 (one-way ANOVA followed by Dunnett *post-hoc* test).

As dissociated cortical neurons randomly form interconnection with neighboring neurons and develop spontaneous network activity via excitatory synaptic transmission in culture, we next pharmacologically examined whether ultrasound-mediated Ca^2+^ transients are induced by the enhancement of network activity (Fig. 2E). Tetrodotoxin (TTX), a selective blocker of voltage-gated sodium channels, inhibited the generation of action potential in individual neurons, and we observed that the toxin treatment elicited no neuronal responses to ultrasound stimuli (Fig. 2F-H). We also tested the effect of CNQX and AP5, potent antagonists against AMPA-type and NMDA-type ionotropic glutamate receptors in post-synapse respectively, on ultrasound dependent neuronal responses. In the presence of both drug mixtures, neuronal responses to ultrasound were sharply suppressed (Fig. 2F-H). Additionally, we confirmed that nifedipine, a selective L-type voltage-gated calcium channel blocker, also significantly reduced ultrasound-induced neuronal responses (Supplemental Fig. 3C). These results suggest that ultrasound-mediated Ca^2+^ transients require the generation of action potentials and functional network activity.

Taken together, our findings indicate that ultrasound-mediated Ca^2+^ transients represent the overall neuronal excitation of network activity arising as a consequence of the biological events involved in ultrasound irradiation.

### TRPC6 is an essential channel for ultrasound-induced neuronal responses

The biophysical effects of ultrasound on living organisms include temperature rise, cavitation, and acoustic radiation pressure. We noted that ultrasound irradiation did not cause a significant change in bath temperature in our experimental condition (data not shown), excluding the possibility that ultrasound-induced neuronal responses are caused by the mechanisms related to a temperature increase. In addition, the ultrasound waves at the frequency of 1 MHz and an intensity of 190 mW/cm^2^, which we used in all experiments, are below the cavitation threshold (23). Thus, we hypothesized that acoustic radiation pressure generates a mechanical force to activate mechanosensitive channels in neurons, which may be an initial trigger for neuronal depolarization and further excitation of network activity. Since cortical and hippocampal neurons express several potential mechanosensitive channels including TRPV1, TRPV2, TRPV4, Piezo1, TRPC1, and TRPC6 in the mammalian brain (17, 24, 25), we next sought to identify the specific channels that predominantly contribute to ultrasound-mediated Ca^2+^ transients using pharmacological blockers (Fig. 3A and E).

**Figure 3.**
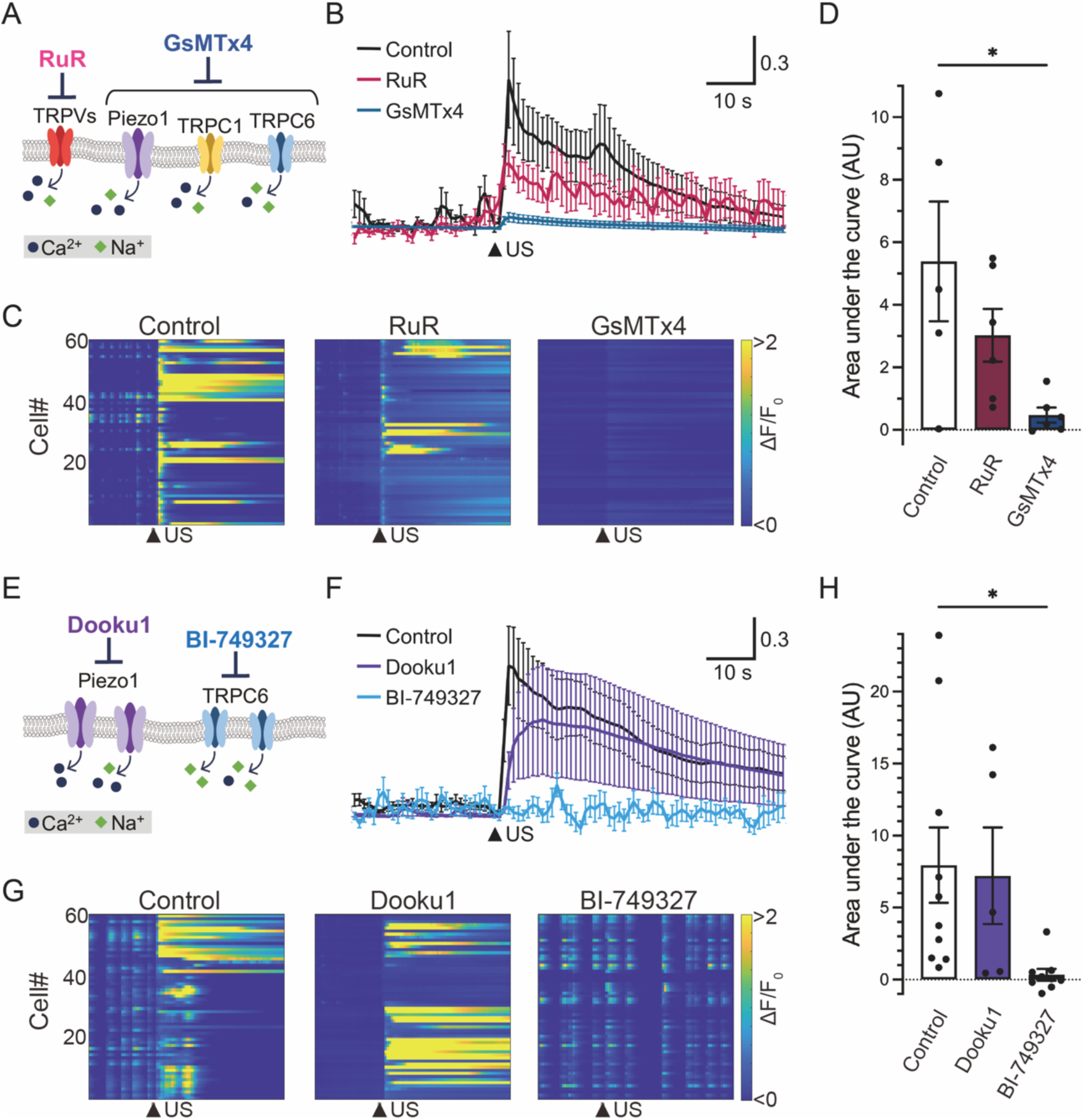
Pharmacological investigation of the involvement of mechanosensitive channels in neuronal responses to ultrasound stimulation. A) A schematic diagram of mechanosensitive receptors and effects of antagonists. Ruthenium red (RuR, 5 µM) was used to block TRPVs channel and GsMTx4 (10 µM) was used to prevent gating Piezo1/2, TRPC1, and TRPC6. B) Averaged traces of neuronal Ca^2+^ transients against ultrasound irradiation under the condition of control (black), RuR-treated (red), or GsMTx4-treated (blue) neurons. Data from five, six, and six independent experiments for control, RuR, and GsMTx4 are described as mean ± SEM, respectively. Arrowhead indicates the time of ultrasound stimulation. C) Heatmap demonstration of normalized fluorescence intensity (ΔF/F_0_) of GCaMP6s in neurons under the condition of control, or the presence of RuR or GsMTx4. Arrowheads indicate the time of ultrasound stimulation. Data from 60 cells in an experiment are plotted. Arrowheads indicate the time of ultrasound stimulation. D) The amplitude of Ca^2+^ transients in responses to ultrasound stimulation is quantified as area under the curve (AU). Bar graph values show mean ± SEM. \**P* < 0.05 (one-way ANOVA followed by Dunnett *post-hoc* test). E) A scheme of the effects of selective antagonists on Piezo1 or TRPC6. Dooku1 (10 µM) and BI-749327 (1 µM) were used to inhibit Piezo1 and TRPC6 activation, respectively. F) Averaged traces of neuronal Ca^2+^ transients against ultrasound irradiation under the condition of control (black), or the presence of Dooku1 (purple) or BI-749327 (light blue). Ten, five, and nine independent experiments for control, Dooku1, and BI-749327 were performed, respectively. Traces show mean ± SEM. G) Heatmap analysis of neuronal responses under the condition of control, Dooku1 treated, or BI-749327 treated neurons. H) Bar graph values of area under the curve in each condition show mean ± SEM. \**P* < 0.05 (one-way ANOVA followed by Dunnett *post-hoc* test).

To narrow down the candidate molecules, we examined the effect of ruthenium red (RuR), the antagonist of the broad spectrum of TRPV channels (Fig. 3B-D). In the presence of RuR, neuronal responses to ultrasound tended to be suppressed, but they did not significantly differ between the control group and drug-treated group. We also tested the effect of GsMTx4, the peptide inhibitor of mechanosensitive ion channels including Piezo1, TRPC1, and TRPC6, on the ultrasound responses of neurons. Remarkably, neurons incubated with GsMTx4 powerfully reduced ultrasound-evoked responses, suggesting Piezo1, TRPC1, and TRPC6 as potential candidates for an ultrasound-responsible molecule (Fig. 3B-D). Thus, we pharmacologically segregated these channel pathways utilizing two additional chemicals BI-749327, a selective TRPC6 antagonist, and Dooku1, a Piezo1 antagonist, on ultrasound-induced neuronal responses. To our surprise, ultrasound mediated Ca^2+^ transients in neurons were almost completely diminished in the presence of BI-749327 (Fig. 3F-H). In contrast, we observed that Dooku1 treatment did not have a significant impact on neuronal responses to ultrasound in our experimental condition (Fig. 3F-H). We also observed neuronal responses induced by the bath administration of Hyp9, a selective TRPC6 agonist, thus confirming the expression of functional TRPC6 channels in cultured cortical neurons (Supplemental Fig. 4).

To further support our findings by genetic experimental evidence further, we performed calcium imaging of cultured cortical neurons isolated from the brain of TRPC6-deficient (TRPC6-KO) mouse embryos (Fig. 4A). Consistent with the results obtained from the pharmacological experiments, TRPC6-KO neurons showed almost no responses to ultrasound irradiation (Fig. 4B-D). Importantly, lentiviral overexpression of the mouse TRPC6 channel in TRPC6-KO neurons restored the neuronal responsive ability against ultrasound stimulation. Taken together, these results strongly indicate that TRPC6 is the sole mechanosensitive ion channel directly activated via ultrasound-driven mechanical force in cultured neurons.

**Figure 4.**
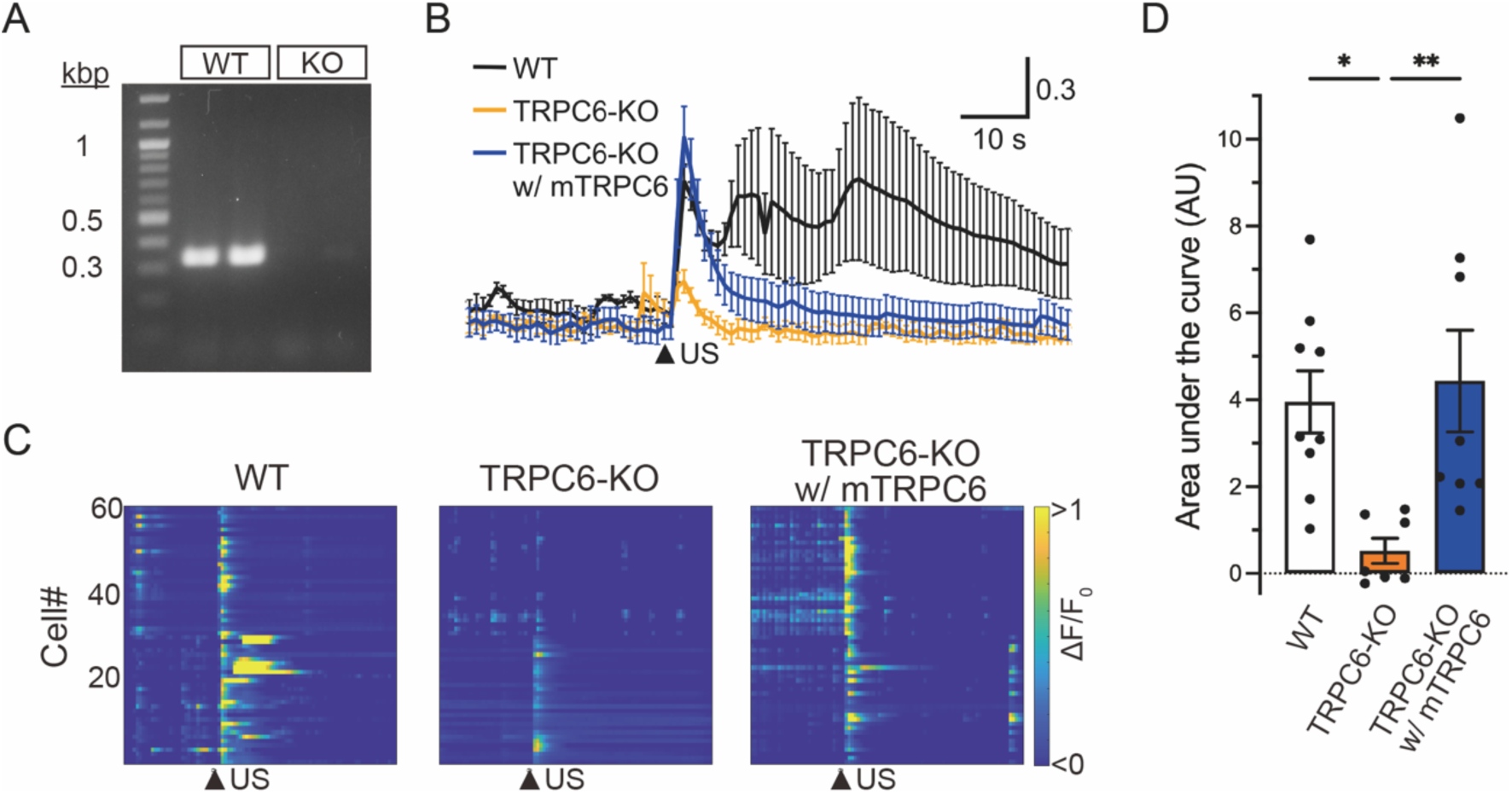
TRPC6-KO neurons lack the response to ultrasound irradiation which can be rescued by overexpression of TRPC6. A) RT-PCR validation of TRPC6 mRNA expression in total RNA extracted from cultured neurons of wild-type (WT) and TRPC6 deficient (KO) mice. Representative gel electrophoresis image from 2 samples each shows PCR products targeting TRPC6 (327 bp). B) Averaged neuronal responses to ultrasound irradiation of WT (black), TRPC6-KO (orange), or TRPC6-KO overexpressing mouse TRPC6-HA (TRPC6-KO w/ mTRPC6) (blue) neurons. Nine, seven, and eight independent experiments for WT, TRPC6-KO, and TRPC6-KO w/ mTRPC6 neurons were performed, respectively. Arrowhead indicates the time of ultrasound stimulation. Traces show mean ± SEM. C) Heatmap demonstration of normalized fluorescence intensity (ΔF/F_0_) of GCaMP6s in WT, TRPC6-KO, and TRPC6-KO w/ mTRPC6. Arrowheads indicate the time of ultrasound stimulation. Data from 60 cells in two experiments are plotted. Arrowheads indicate the time of ultrasound stimulation. D) Bar graph values of area under the curve in each condition show mean ± SEM. \**P* < 0.05, \*\**P* < 0.01 (one-way ANOVA followed by Dunnett *post-hoc* test).

### TRPC6 is essential for ultrasound neuromodulation in mouse brain

Finally, to investigate whether TRPC6 has a critical role in ultrasound-induced neuromodulation in the brain, we performed *in vivo* electrophysiological recordings of population neural activities in the cerebral cortex of anesthetized mice with a pharmacological intervention of TRPC6 (Fig. 5A, B). Transcranial ultrasound irradiation of 1 MHz in 100 Hz PRF and 50% duty cycle, the same parameter as in our *in vitro* experiments, significantly increased population neural activities in the cerebral cortex in an intensity-dependent manner (Fig. 5C, E; P < 0.0001, Χ^2^ test). Notably, consistent with the results of the *in vitro* experiments, the intracerebroventricular administration of BI-749327, an antagonist of TRPC6, significantly reduced activation of cortical neurons induced by transcranial ultrasound irradiation at the intensities of 130, 210, and 350 mW/cm^2^ (Fig 5D, E). Administration of the vehicle did not reduce the responses (data not shown). These results indicate that TRPC6 is a crucial molecule, consistently, in ultrasound neuromodulation both *in vitro* and *in vivo*.

**Figure 5.**
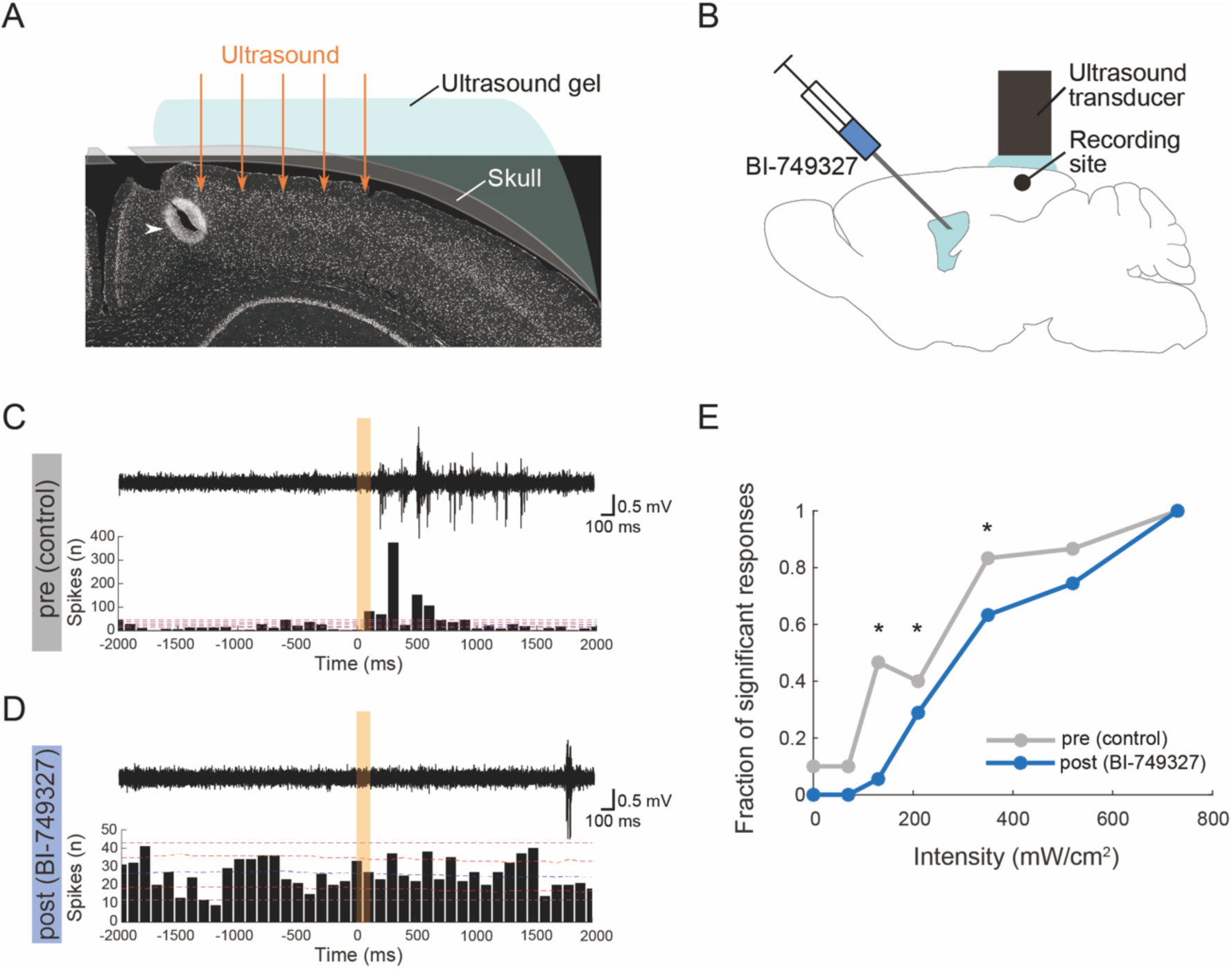
TRPC6 mediates ultrasound-induced neuronal activation *in vivo*. A, B) Schematic diagrams of *in vivo* multi-unit recordings with a TRPC6 blocker, BI-749327. A) Ultrasound was transcranially irradiated to the cerebral cortex of mice, and a tungsten microelectrode was inserted for recordings was inserted through a cranial window. A white arrowhead indicates a representative recording site labeled with an electrical lesion after recordings. B) BI-749327 was intracerebroventricularly administered 30 min before post recordings. C, D) Representative in vivo multi-unit recordings around US irradiations before (C) and after (D) BI-749327 administration. Top: high-pass filtered extracellular voltage traces; Bottom: peri-stimulus time histograms (PSTHs). Orange patches represent ultrasound irradiation. Dashed pink, red and blue lines in each PSTH represent global significant levels, local significant levels, and mean counts calculated with 1,000 randomly generated datasets of timestamps of population neural activities with time jittering, respectively. The significance of each recording was determined by whether PSTH goes above or down across the global significant levels. E) Fraction of significantly activated population responses by US irradiations of several intensities. Significance on each stimulus intensity was determined by Χ^2^ test on a two-by-two (significance x control or BI-749327) frequency table. n = 73, 80 sessions from three mice for pre and post recordings, respectively. \**P* < 0.05.

## Discussion

In the present study, we established experimental systems to assess neuronal activities evoked by ultrasound irradiation compatible with both cortical neuronal culture and living mouse brain, and we examined the detailed molecular and cellular bases of ultrasound neuromodulation in mammalian neurons. A line of evidence has indicated that the acoustic pressure of ultrasound mechanically impacts neurons, leading to rapid network excitation via the generation of action potentials and synaptic transmission. Most importantly, we successfully identified TRPC6, a mechanosensitive cation channel, as a pivotal biosensor that can detect the mechanical pressure of ultrasound and trigger the sequential biological events of intrinsic neuronal responses. Since ultrasound neuromodulation has attracted increasing attention as an innovative technology to control brain circuits with high spatiotemporal precision, our findings offer crucial insight into the future feasibility of safe and fine-tunable ultrasound neuromodulation supported by scientific evidence, which may eventually contribute to a wide range of neuroscience research fields and disease therapies in humans.

TRPC6 is one of the non-selective cation channels widely expressed in various organs including the heart, kidney, and brain (25–27), and it is involved in the regulation of vascular smooth muscle contractility and arterial blood pressure (18). In agreement with previous studies characterizing neuronal expression and dendritic localization of TRPC6 in mouse forebrain (24, 28, 29), we distinctly detected endogenous TRPC6 mRNA in cultured mouse cortical neurons. While several research groups have demonstrated that TRPC6 acts as a mechanosensor in cardiomyocytes (30) and podocytes (31) to sense hypoosmotic stretch or indentation of the plasma membrane, there have been no such reports on neuronal cells of the central nervous system. Therefore, the present study provides the first evidence that TRPC6 is an intrinsic mechanosensitive channel in cortical neurons essential for ultrasound neuromodulation both *in vitro* and *in vivo*.

We propose a hypothetical model in which ultrasound-induced Ca^2+^ transient in neurons arises as a consequence of the generation of action potentials and neuronal network activity triggered by TRPC6 activation (Fig. 6). In this model, TRPC6 may act as a biosensor to detect the mechanical distortion of the lipid bilayer structure on the plasma membrane caused by the acoustic pressure of ultrasound. Since spontaneous Ca^2+^ oscillation of neuronal culture was relatively intact in the presence of selective TRPC6 antagonist or the genetic deficiency of TRPC6 (data not shown), we assume that TRPC6 may not have an essential role in spontaneous network activity. In addition, as TRPC6 has lower calcium ion permeability than other TRP channels such as TRPVs (32, 33), the opening of the TRPC6 channel would preferentially contribute to initial membrane depolarization through sodium influx followed by a generation of the action potential in the small neuronal population. Consequently, network excitation with robust Ca^2+^ transient can be amplified through excitatory glutamatergic synaptic transmission (Fig. 6). This sequential biological process presumably takes a couples of hundred milliseconds, and it can explain the delayed onset of neuronal response after ultrasound stimulation (Fig.5).

**Figure 6.**
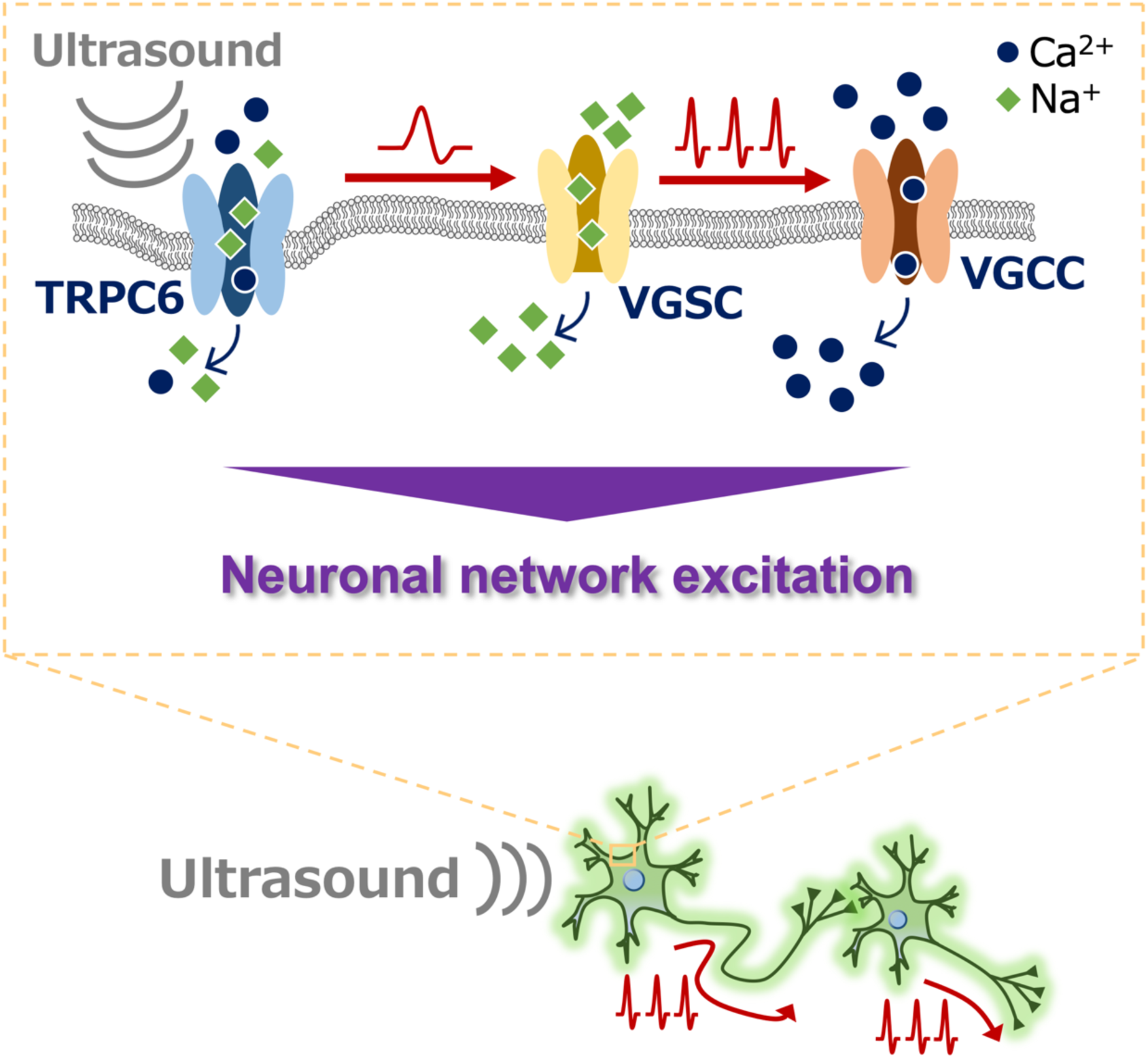
Proposed model for the mechanism of ultrasound neuromodulation via TRPC6 activation. Ultrasound irradiation potentially distorts the cell membrane by acoustic pressure, which triggers the opening of a mechanosensitive TRPC6 channel followed by membrane depolarization in cultured cortical neurons. The generation of action potentials induced by the opening of voltage-gated sodium channels (VGSC) following the TRPC6-mediated depolarization may further elicit extracellular Ca^2+^ influx via voltage-gated calcium channels (VGCC), resulting in excitation of the neuronal network.

*In vitro* culture assay has superior compatibility with pharmacology and genetics, and serves as a powerful experimental platform for screening essential factors involved in ultrasound neuromodulation. Several research groups have independently reported that multiple mechanosensitive ion channels such as TRPC1, TRPP1/2, TRPM4 (16), Piezo1 (14), and N-Methyl-D-aspartate receptor (34) are essential in neuronal responses to ultrasound irradiation in mammalian cortical and hippocampal neurons. We consider that these controversial outcomes from several laboratories including from our group may be due to the huge variability among the experimental configurations of each laboratory. For instance, while neurons in our experimental system uniformly received direct ultrasound stimulation from above in our experimental system, other groups stimulated neurons by indirect sonication through thin film or glass coverslips from the bottom side of the recording chamber (14, 16). In addition, multiple parameters of acoustic waves (i.e. flat or focused, frequency, intensity, duty cycle, duration, radiation angle, etc) complicate the major bioeffect of ultrasound that may impact the specificity and efficacy of the activation or inactivation of target molecules in neurons (9). In this regard, standardization of experimental systems with optimized ultrasound parameters would be necessary to establish sufficient consensus and comprehensive knowledge for ultrasound neuromodulation.

Another important aspect of our findings is that the inhibition of the TRPC6 channel consistently diminishes the neuronal response to ultrasound both *in vitro* and *in vivo*. Although immortalized cell lines are relatively easy to handle and maintain, they cannot develop network activity with synaptic connections and would not be an appropriate model for investigating brain function. In contrast, our primary neuronal culture demonstrates spontaneous network oscillation mediated by synaptic transmission and well represents the physiological brain state of living animals. We could reproduce reliable results indicating that TRPC6 is consistently essential for ultrasound neuromodulation both *in vitro* and *in vivo*, and therefore it is most likely that our findings reflect the biological phenomena observed in the human brain in clinical studies. Nonetheless, we cannot rule out the possible involvement of other molecules, the biophysical properties of the plasma membrane, and cellular and network mechanisms. For example, we still observed a high probability of neuronal responses to ultrasound even in the presence of TRPC6 blocker when increased irradiation intensity (Fig. 5). This may be due to insufficient permeability of the drug from the ventricles into the cortical parenchyma, or it may also be due to other mechanisms such as reorganization of organelles and cytoskeletons or contribution of other mechanosensitive channels (15, 35). Since diverse cell types such as neuronal subtypes (36), astrocytes (35), and microglia (37) may respond to ultrasound differently, further investigation would be required to fully understand ultrasound neuromodulation.

Current technical advances in a variety of neuromodulation modalities have tremendously contributed to the progress of neuroscience. The most widely used neuromodulation techniques are optogenetics and chemogenetics in animal studies. Optogenetics enables accurate control of neuronal activities by light illumination with high spatiotemporal resolution but requires invasive surgical implantation of optical fiber. In contrast, chemogenetics achieves robust alteration of basal neuronal activities by systemic drug administration in a less invasive manner; but chemogenetics lacks spatial or temporal resolution, which are both important for selective and effective intervention of the neuronal network *in vivo* (38). Importantly, both optogenetics and chemogenetics require gene transduction in targeted brain circuits by local virus injection or DNA electroporation, and therefore these approaches are not easily applicable to the human brain. Compared to these technologies, ultrasound has superior advantages in terms of its non-invasiveness and remarkable tissue permeability. In addition, the transcranial ultrasound wave can be focused to concentrate its acoustic energy in the target region, offering unique applications to modifying specific brain circuits with high spatiotemporal regulation. Indeed, a recent study has demonstrated that transcranial focused ultrasound stimulates the hypothalamus preoptic area via TRPM2 channels and causes torpor-like hypothermic and hypometabolic states in rodents (39), promising the feasibility of ultrasound neuromodulation of deep brain circuits in living animals. Furthermore, growing evidence has highlighted the conceptual advance of sonogenetics, and an approach to utilizing genetically encoded ultrasound-responsive mechanosensitive molecules for control of neural activity of genetically defined neuronal population, as a next-generation neuromodulation technology (9, 22, 40). Although the applicability of multiple mechanosensitive channels, including mechanosensitive channel of large conductance (MscL, a thoroughly analyzed force sensor of *Escherichia coli*) (41–43), Piezo1 (44) and TRPA1 (35, 45) for sonogenetics has been reported in several laboratories, it is still in its beginning phase and the topic is beyond the scope of the present study. Additional effort would be necessary in order to establish sonogenetics as a mature technical platform in neuroscience.

In conclusion, we found that TRPC6 is an essential intrinsic mediator of ultrasound neuromodulation in cortical neurons of the mammalian brain. Since it has been reported that transcranial ultrasound neuromodulation ameliorates the brain pathologies and behavior abnormalities of animal models for neurodegenerative disorders, such as Parkinson’s Disease and Alzheimer’s Disease (46, 47), TRPC6 may contribute to the beneficial mechanism in ultrasound neuromodulation and can also be an attractive pharmacological target for disease-modifying therapies. Our findings provide new insight for the encouragement of further investment in ultrasound neuromodulation, and the application of this modulatory technology to animal models will provide *in vivo* evidence for pivotal roles played by the discovered mechanical processes.

## Acknowledgements

We thank Dr. Nobuki Kudo for the helpful discussion and technical suggestion and Biology of Non-domain Biopolymer Grant-in-Aid for Transformative Research Area (21A304) for access to a confocal microscopy. We also thank all laboratory members for their discussions and continuous encouragement. This work was supported in part by Grants-in-Aid for Brain Mapping by Integrated Neurotechnologies for Disease Studies (Brain/MINDS; 18dm0207018 (M.H.), 19dm0207072 (M.H.), and 20dm0207072 (M.H.)), JST Grant Number JPMJMS2024 (M.H.), JSPS KAKENHI Grant Numbers JP21K07580 (M.S.), JP23K14312 (Y.M.), JP23H02790 (Y.T.), and JP23K18250 (Y.T.), QST President’s Strategic Grant Exploratory Research (Y.M.), and AMED under Grant Number JP24gm6510013 (M.S.), JP24gm6510015 (Y.T.), JP24zf0127004 (Y.T.), JP21zf0127004 (M.H.), and 22zf0127004s1002 (M.H.).

## Author contributions

Y.M., M.S., and M.H. conceived the study; Y.M. and M.Y. constructed plasmid DNA and generated recombinant virus vectors; Y.M., M.Y., and M.S. performed biochemistry, live imaging, histology, and data analysis. T.S. carried out RT-PCR analysis; S.T recovered TRPC6-KO mice from cryopreserved fertilized eggs; K.Y. and Y.T. performed in vivo electrophysiology; Y.M, M.S., K.Y., Y.T., and M.H. wrote the manuscript.

---

**Supplemental Figure 1.**
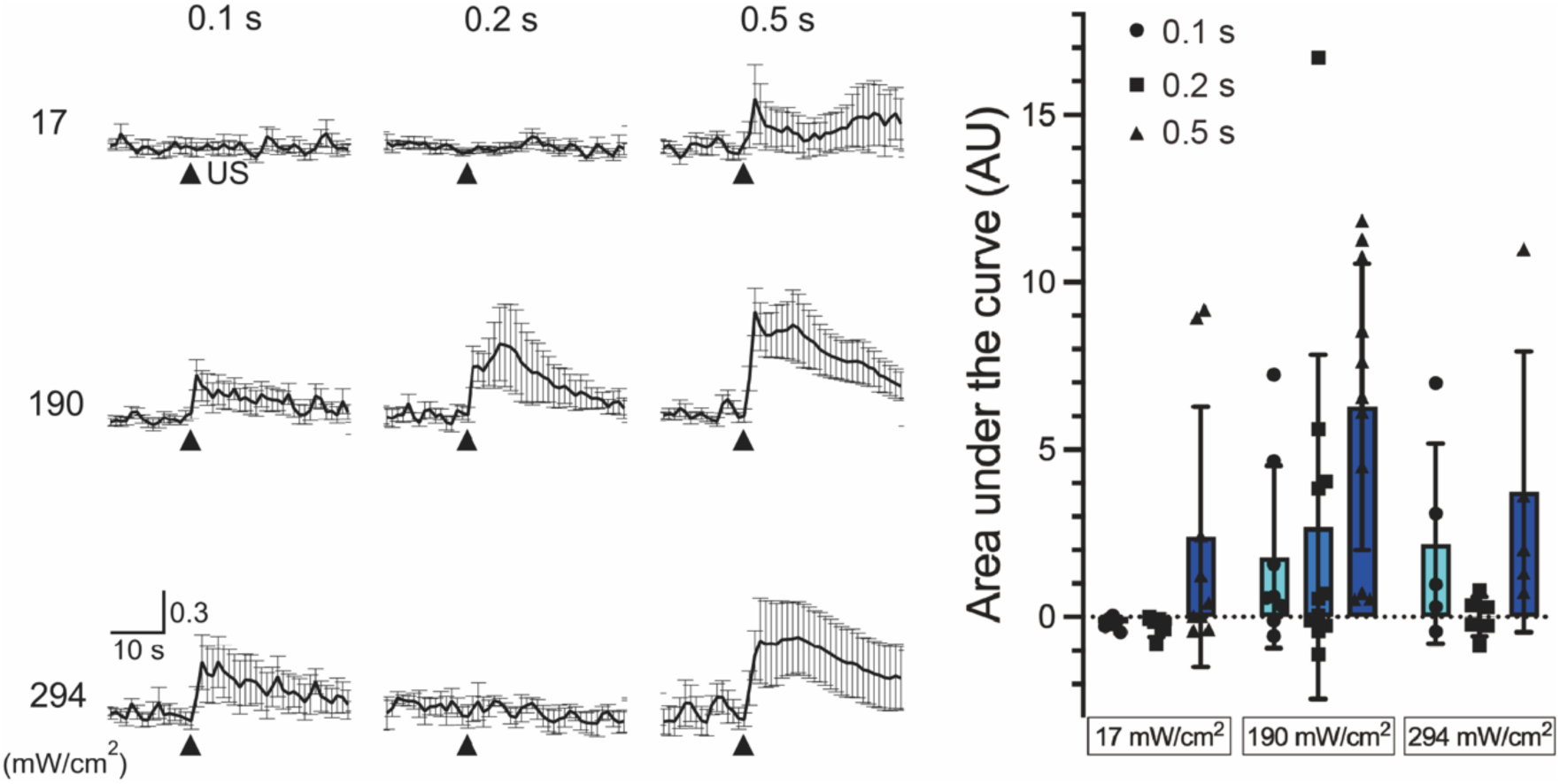
Ultrasound-induced neuronal responses and quantification of the responses as function of ultrasound intensity and stimulus duration. Stimulus intensities of 17, 190 and 294 mW/cm^2^ were used, and they were stimulated for 0.1 (n = 6, 8, 5), 0.2 (n = 5, 11, 6), and 0.5 (n = 9, 11, 5) seconds, respectively. Arrowheads indicate the time point of ultrasound stimulation. The amplitude of Ca^2+^ transients in responses to ultrasound stimulations is quantified as area under the curve (AU). Traces and bar graph values show mean ± SEM.

**Supplemental Figure 2.**
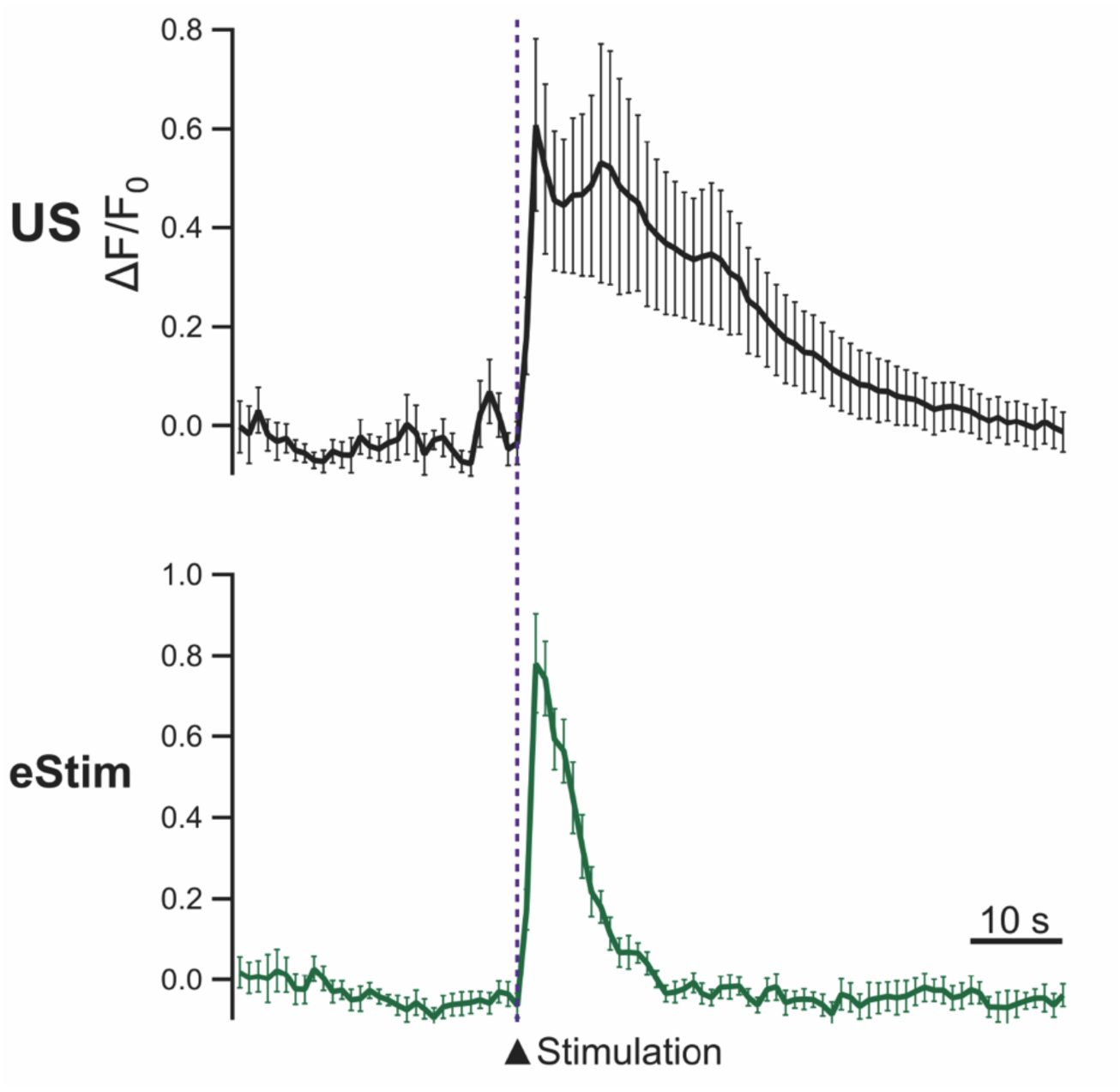
Comparison of ultrasound-induced and electrical stimulation-induced neuronal calcium responses. Neuronal calcium responses against ultrasound irradiation (US) or electrical stimulation (eStim) of cultured neurons. Eleven and ten independent experiments for US and eStim were performed, respectively. Traces values show mean ± SEM.

**Supplemental Figure 3.**
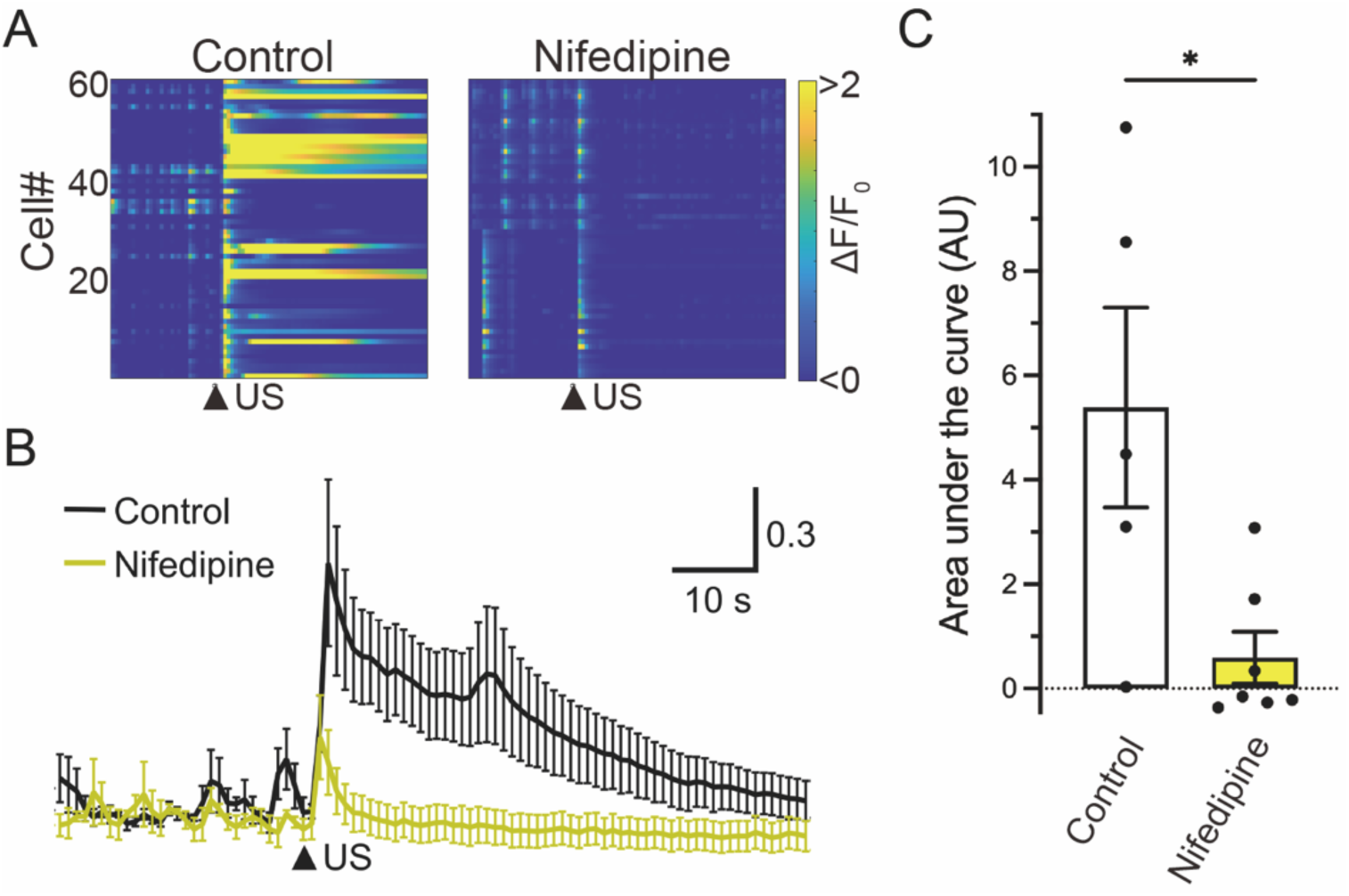
Neuronal calcium responses induced by ultrasound irradiation in the presence of nifedipine. A) Heatmap demonstration of normalized fluorescence intensity (ΔF/F_0_) of GCaMP6s in neurons under the condition of control or the presence of nifedipine (100 µM). Nifedipine is used to block L-type voltage-gated calcium channels. Arrowheads indicate the time of ultrasound stimulation. B) Averaged traces of neuronal Ca^2+^ transients against ultrasound irradiation under the condition of control (black), or the presence of nifedipine (yellow). Data from five and seven independent experiments for control and nifedipine are described by mean ± SEM, respectively. C) Bar graph values of area under the curve in each condition show mean ± SEM. \**P* < 0.05 (one-way ANOVA followed by Dunnett *post-hoc* test).

**Supplemental Figure 4.**
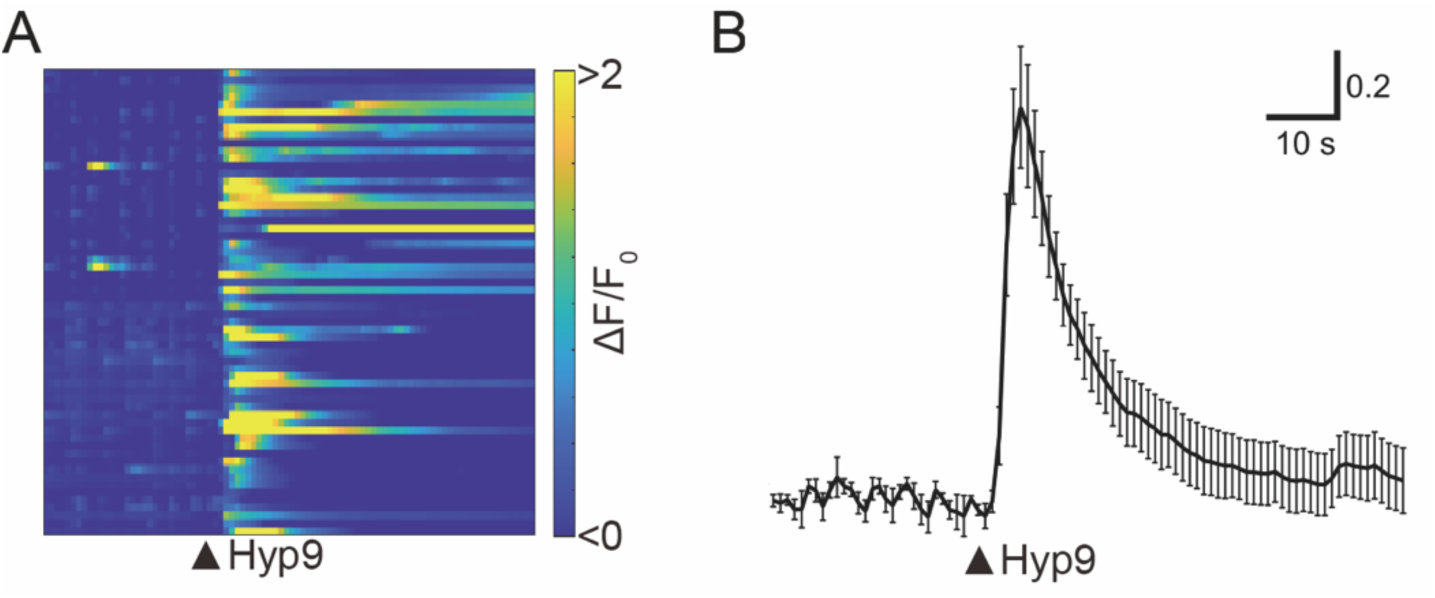
Neuronal responses to a selective TRPC6 agonist, Hyp9. A) Heatmap demonstration of normalized fluorescence intensity (ΔF/F_0_) of neuronal responses when Hyp9 (10 µM) is applied. Arrowheads indicate the time of Hyp9 application. Data from 60 cells in two experiments are plotted. B) Averaged neuronal response to Hyp9 (n = 3).

